# Burn Wound Excision Drives Acute Hypermetabolism in Mice with Scald Burns

**DOI:** 10.64898/2026.10.26.761985

**Authors:** Jaycelyn S Hall, Abigail Barlow, Mary C Barre, Lillie D Treas, Richard Yeager, Esther Teo, Meagan S Kingren, Craig Porter

## Abstract

Hypermetabolism (elevated resting energy expenditure) is a hallmark of the pathophysiological response to burns. The contribution of wound excision to acute burn-induced hypermetabolism remains unclear. Importantly, while burn wound excision is a standard component of clinical burn care, it is not typically included in rodent models of burn injury. Here, we tested the hypothesis that burn wound excision drives the acute hypermetabolic response to burn trauma.

C57BL/6J mice (n=12-18 per group; ∼6-weeks old; 50% male) were acclimated to thermoneutral housing (30°C) for 6-weeks. All mice underwent baseline body composition scans and metabolic phenotyping to determine total (TEE) and resting energy expenditure (REE). Mice were then randomized to either sham (control), scald burn (burn), wound excision (excision), or scald burn followed by immediate wound excision ( burn+excision) groups. Burns and wound excisions involved ∼15% of the body surface area. Mice then recovered for 14 days while individually housed in metabolic phenotyping cages. At day 14 post-injury, mice underwent a second body composition scan prior to euthanasia.

There were no fatalities in the control or burn groups, whereas 23% (n=7) and 34% (n=11) of mice in the excision and burn+excision groups died or were euthanized during the 14-day recovery period. Male and female mice exhibited broadly similar responses to injury in terms of REE and TEE. Compared to control (3.59±0.56 kcal/day), REE was greater in burn (5.29±1.03 kcal/day), excision (7.25±0.84 kcal/day), and burn+excision groups (8.48 ± 0.91 kcal/day) during the first week post-injury (P<0.001 for all). Accordingly, the magnitude of hypermetabolism was greater in the excision (+102%) and burn+excision groups (+136%) versus burn alone (+47%) and control. Compared to control mice, wound excision (alone or following burns) resulted in a greater acute loss in body mass than burn injury alone. However, this cachectic response was more sustained in male mice compared to females. Similarly, wound excision (alone or following burns) resulted in greater bone mineral content and density loss compared to sham and burn alone in both sexes.

Wound excision is a key contributor to the hypermetabolic and cachectic response to burn trauma in mice. We posit that rodent studies aiming to recapitulate burn-induced hypermetabolism and cachexia should include burn wound excision to better model the clinical sequelae to burns.

## Introduction

Hypermetabolism, an increase in whole body resting energy expenditure (REE), is a hallmark of the metabolic stress response to burn injury^1–3^ and is associated with cachexia, delayed healing, and long-term morbidity^4,5^. From a biochemical perspective, increased ATP synthesis to support wound healing, the acute phase response, and intermediary metabolism^6^, as well as recruitment of mitochondrial proton leak to offset heat loss due to a compromised skin barrier^7^, underlie burn-induced hypermetabolism. While early burn wound excision has been shown to aid recovery and prevent complications such as infection in patients^8^, it is likely that excision of burn wounds contributes to acute burn-induced hypermetabolism due to evaporative heat loss and a heightened demand for substrates and ATP to support wound healing.

Patient data demonstrate a relationship between burn wound size and evaporative heat loss^9–11^, which can be mitigated to a degree by occlusive wound dressing^12,13^ and ambient temperature ^7^. Furthermore, in patients that underwent early versus delayed total burn wound excision, REE markedly increased post wound excision^8^. These observations have important implications for preclinical burn research, as they suggest that to accurately model the acute hypermetabolic response to burns in rodents, wound excision should be incorporated. Accordingly, the current study was designed to test the hypothesis that burn wound excision amplifies the acute hypermetabolic response to burn trauma. In mice acclimated to thermoneutral housing temperatures prior to experimental procedures, we comprehensive ly characterized the hypermetabolic-catabolic responses to burn injury alone, wound excision alone, and combined burn injury with subsequent excision of burn wounds. Our data suggest that under thermoneutral housing conditions, wound excision is a key driver of the acute hypermetabolic catabolic stress response to burn trauma.

## Methods

### Animals

The Institutional Animal Care and Use Committee (IACUC) of the University of Arkansas for Medical Sciences reviewed and approved all the animal procedures and care methods. Young (∼6 weeks old) male and female C57BL/6J (Jackson Laboratories, Bar Harbor, ME) mice were used in this study. Upon arrival, animals were group housed and acclimated to a thermoneutral housing temperature (30°C) for at least ∼4-weeks on a reverse light cycle (light 7:00 pm-7:00 am) with ad libitum access to water and a standard chow diet [TD.95092 (18.8% protein, 17.2% kcal fat, 63.9% kcal carbohydrate, 3.8 kcal/g), Envigo Teklad Diets, Madison, WI]. Mice were then singly housed in metabolic phenotyping cages for a baseline determination of energy expenditure for one week. At the end of the week, animals underwent a dual energy x-ray absorptiometry (DEXA) scan.

An overview of the study protocol is shown in **Figure 1**. Animals were randomized based on their average daily energy expenditure and body masses to either sham (control), scald burn (burn), wound excision (excision), or scald burn followed by immediate wound excision (burn+excision) groups^14^. All mice were administered ∼3 mg/kg extended-release buprenorphine (Covetrus) subcutaneously ∼90 minutes prior to sham/excision/burn procedures. Mice were then anesthetized using isoflurane ( ∼2.5%), and the dorsum and flanks were shaved using electrical clippers, after which up to 1 mL of lactated ringers was injected subcutaneously above the spine. Mice were then placed in a customized cast that exposed either 8.6 cm^2^ (males) or 6.5 cm^2^ (females) of the dorsum. For mice in the excision and burn+excision groups, a wound outline was drawn on the dorsum around the outside of the cast. Still in the cast, mice in the burn and burn+excision groups, mice were then submerged in ∼95°C water for ∼10s while under isoflurane anesthesia. Mice randomized to excision or burn+excision were transferred to a sterile surgical table and remained under isoflurane anesthesia. The entire dorsum was cleaned three times in an iodine solution and then three times with an alcohol solution. The marked wound was then dissected down to the underlying muscle using surgical scissors. Wounds were then coated in a petroleum jelly based topical antibiotic (Neosporin). Mice randomized to excision alone underwent the same excision procedure (without prior burn injury).

**Figure 1:**
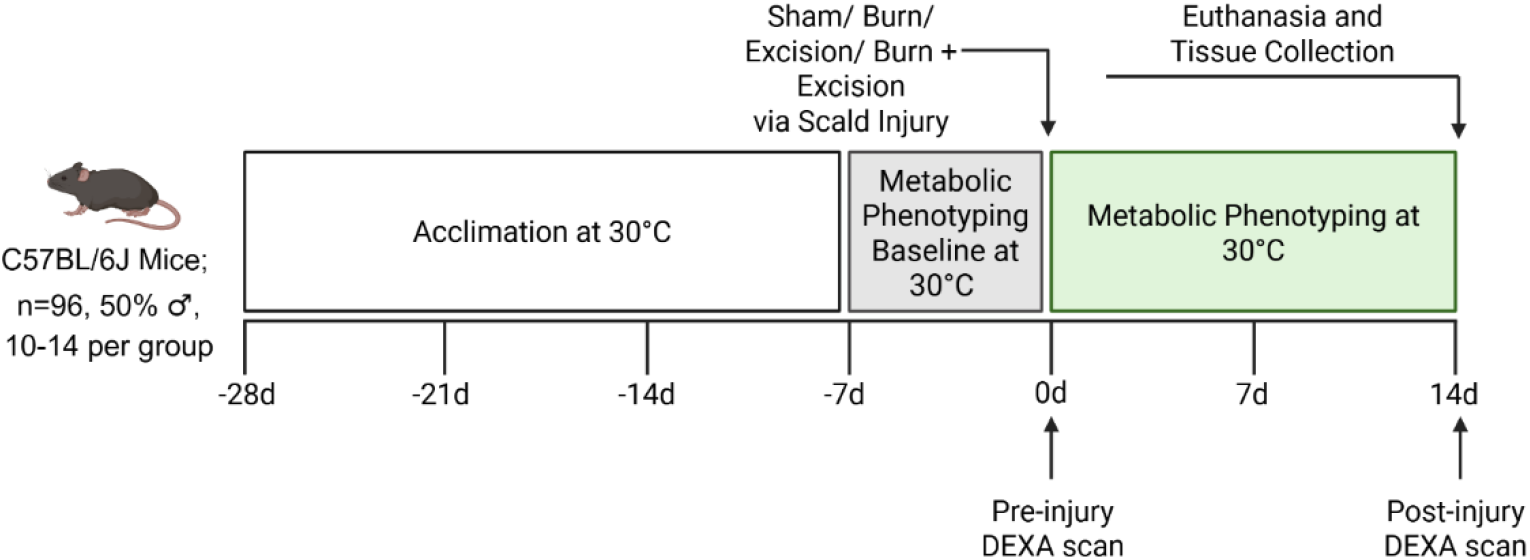
C56BL/6J mice were acclimated at 30°C for 3 weeks. Mice were then singly housed in metabolic phenotyping cages for an additional ∼1 week at 30°C to establish a metabolic baseline. Then, mice were randomly assigned to a sham, burn, excision, or burn+excision groups, blocking for baseline metabolism and body weight (sham/burn: n=10 per group/sex. Excision/burn+excision: n=13-14/group/sex). Mice received a pre-injury DEXA pre-procedure and a post-injury DEXA scan on the day of euthanasia.

All animals were resuscitated with up to 1 mL of lactated Ringers’ buffer (i.p.) and monitored until fully ambulatory following anesthesia. Sham animals underwent the same procedure (buprenorphine administration, dorsum and flank shaving, and fluid resuscitation), but without burn/excision/burn+excision procedures. All mice were singly housed at 30°C in metabolic phenotyping cages for 14 days post procedure. Mice had unrestricted access to water and diet during this 14-day period. Up to day 7 post procedure, all mice also had access to ∼5 grams of an H_2_O gel-pack (ClearH_2_O) and moist diet pellets (∼12 grams) placed on their cage floor to aid hydration and feeding. Prior to euthanasia on day 14, all mice underwent a second DEXA body scan. Mice were euthanized in a rising concentration of CO_2_ and blood and tissues were collected.

### Estimation of Body Surface Area and Body Composition

To calculate relative burn or excision size, we first calculated total body surface area (TBSA) using the Meeh constant of 6.578 (*k*)^15^ for C57BL/6J mice using the formula: *TBSA* = *k x* mass^0.677^. This resulted in average TBSA for male mice of 61.0 cm^2^ and for females of 48.1 cm^2^ (**Tables 1, 2**). The wound size used for both the burn and excision wound was 8.6 cm^2^ for male and 6.5 cm^2^ for female mice.

**Table 1.**
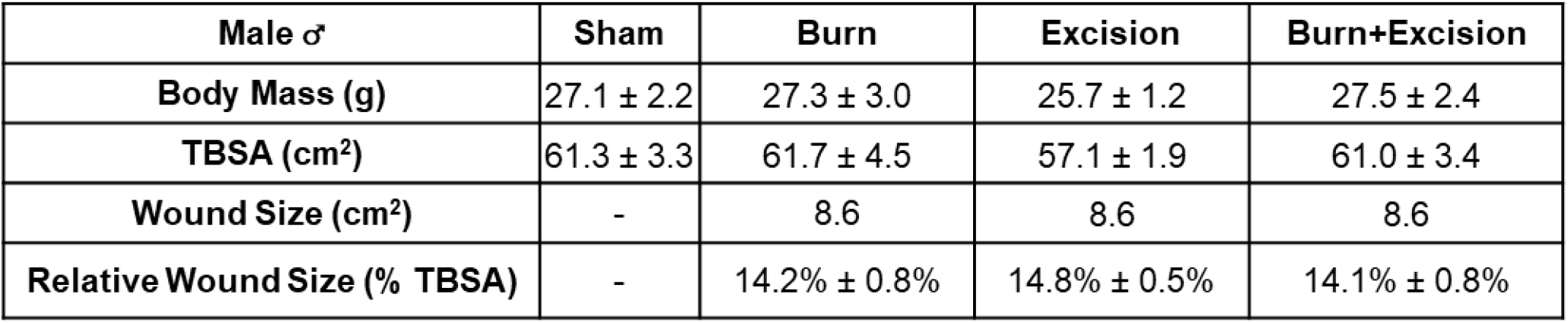
Body mass, estimated total body surface area (TBSA), absolute wound size, and relative wound size for male mice. Values are group means ± SD. n=9-11 for per group based on survivors.

**Table 2.**
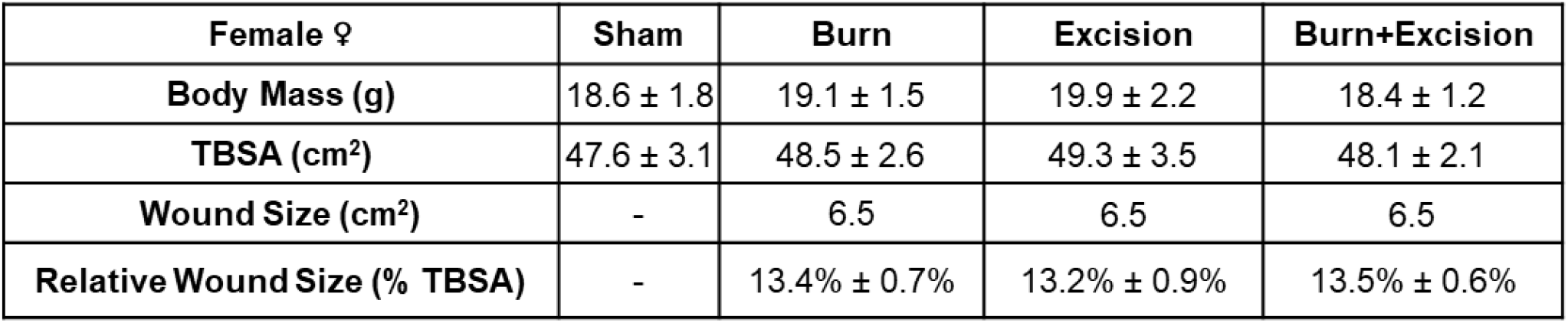
Body mass, estimated total body surface area (TBSA), absolute wound size, and relative wound size for female mice. Values are group means ± SD. n=9-11 per group based on survivors.

Body composition was measured by dual energy x-ray absorptiometry (DEXA) (iNSiGHT VET DXA; Osteosys, Seoul, South Korea). Mice were briefly anesthetized in ∼2.5% isoflurane and placed on the scanner bed in a prone position. Body composition baselines were determined during the light phase prior to procedures (Day 0) and again on the day of euthanasia (Day 14).

### Metabolic and Behavioral Phenotyping

For ∼1 week prior to procedures and during the 14-day recovery period, 64 (7 per group for sham/burn, 9 per group for excision/burn+excision) mice underwent metabolic and behavioral phenotyping. Mice were individually housed in specialized cages capable of continuously measuring VO_2_, VCO_2_, and water vapor production (Promethion Core, Sable Systems International, Las Vegas, NV) (**Supplemental Fig 1**). Feeding, drinking, and ambulation were also continuously measured. During the first week post-injury, mice were supplied with supplemental diet and H_2_O gel (∼12 and ∼5 grams, respectively) in their cages, though amount consumed was not measured. Due to the supplemental hydration supplied to injured animals, corrections were made to the water vapor production values. Supplemental hydration was placed in empty cages to determine the contribution of evaporation of H2O gel to water vapor production rate measure by the metabolic cage system. Based on these values (∼0.28 ml/hr), the water vapor production caused by supplemental hydration was subtracted from the measured values in cages (**Supplemental Fig 2**). Animals were randomized to their position within specialized environmental cabinets designed to control ambient temperature and humidity during metabolic and behavioral phenotyping. Cabinet temperature was set to 30°C for all phenotyping experiments.

Raw data was processed using macros to distill data into hourly averages/totals and circadian reports of phenotyping data for each 12-h light cycle (Sable Systems International, Las Vegas, NV). Energy expenditure rates were calculated from VO_2_ and VCO_2_ using the Weir equation^16,17^. Daily total energy expenditure was calculated as the sum of the average hourly rate of energy expenditure (kcal/hr) for both the light and dark cycle. Daily resting energy expenditure was calculated from the energy expenditure rate during the 30-minute period with the lowest energy expenditure (kcal/h) during the light cycle multiplied by 24 to express REE in kcal/day. Evaporative heat loss was reported as VH_2_O (mL/min) as a measure of whole body water vapor loss. Locomotion was reported as pedestrian meters, which was the measu rement of cage ambulation that only included gross movements. Daily food and water intake were reported as cumulative totals over the 14 day recovery period, and banked per day and per week for each group.

### Tissue Collection

Immediately following euthanasia, tissues were collected and weighed for analysis. Blood was collected into a K+EDTA coated syringe and underwent a complete blood cell count with differential (Sysmex 330). Major organs (heart, liver, kidneys, and spleen) were carefully dissected and weighed, as were the inguinal white adipose tissue (iWAT) and gonadal white adipose tissue (gWAT) depots.

### Statistical Analyses

A priori sample size estimation: Our primary outcome measure for this study was REE. Preliminary data from mice transitioned from an ambient temperature of 24°C to 30°C used in an *a priori* sample size estimation. Assuming a type I error rate of 0.05, we estimated that to detect a physiologically relevant difference in REE that n=8 (power of 0.9) animals would be required per group. To promote rigor within our study design, we determined the impact of sex on the metabolic response to burns. We studied a total of n=20 mice (n=10 males, n=10 females) per group in the sham and burn groups. For excision and burn+excision, we studied n=28 mice total per group (n=14 male, n=14 female) to compensate for mortality within these groups.

Randomization: Within each sex and batch, mice were block randomized by their average daily energy expenditure and by body mass to treatment group (sham, excision, burn or burn+excision) using RandoMice^14^. Within each batch of 32 mice, animals were then randomly assigned to one of the 16 cage positions within metabolic cage systems.

Post hoc data analysis: Data are presented as means±SEM unless stated otherwise. To determine main effects (i.e., injury and sex) and potential significant interactions between time and injury, a Two-Way ANOVA was performed. Multiple comparisons between experimental conditions over time were adjusted for using Tukey’s Test. Comparisons between one time point utilized a One-Way ANOVA and adjusted for using Tukey’s test. Statistical analyses were carried out using GraphPad Prism version 11 (GraphPad Software, LLC, San Diego, CA). Statistical significance was set at P≤0.05.

## Results

### Absolute and Relative Wound Size

Body mass and estimated total body surface areas (TBSA) for male and female mice are presented in **Table 1** and **Table 2**, respectively. Absolute wound sizes for mice in the burn, excision, and burn+excision groups were ∼8.6 cm^2^ for male mice and ∼6.5 cm^2^ for female mice. In male mice, the calculated relative wound sizes were 14.4% for the burn group, 14.6% for the excision group, and 14.4% TBSA for burn+excision group (**Table 1**). For female mice, the calculated relative wound size was 13.8% TBSA for the burn group, 12.9% TBSA for excision group, and 13.3% TBSA for burn+excision group (**Table 2**).

### Mortality

There was no mortality in the sham or burn groups. In male mice, 19% of the excision group and 25% of the burn+excision group died or were euthanized within the first 48 hr. post injury (**Fig 2A**). In female mice, 27% of the excision group and 31% of the burn+excision group died or were euthanized, typically within the first 48 hr. post injury (**Fig 2B**).

**Figure 2:**
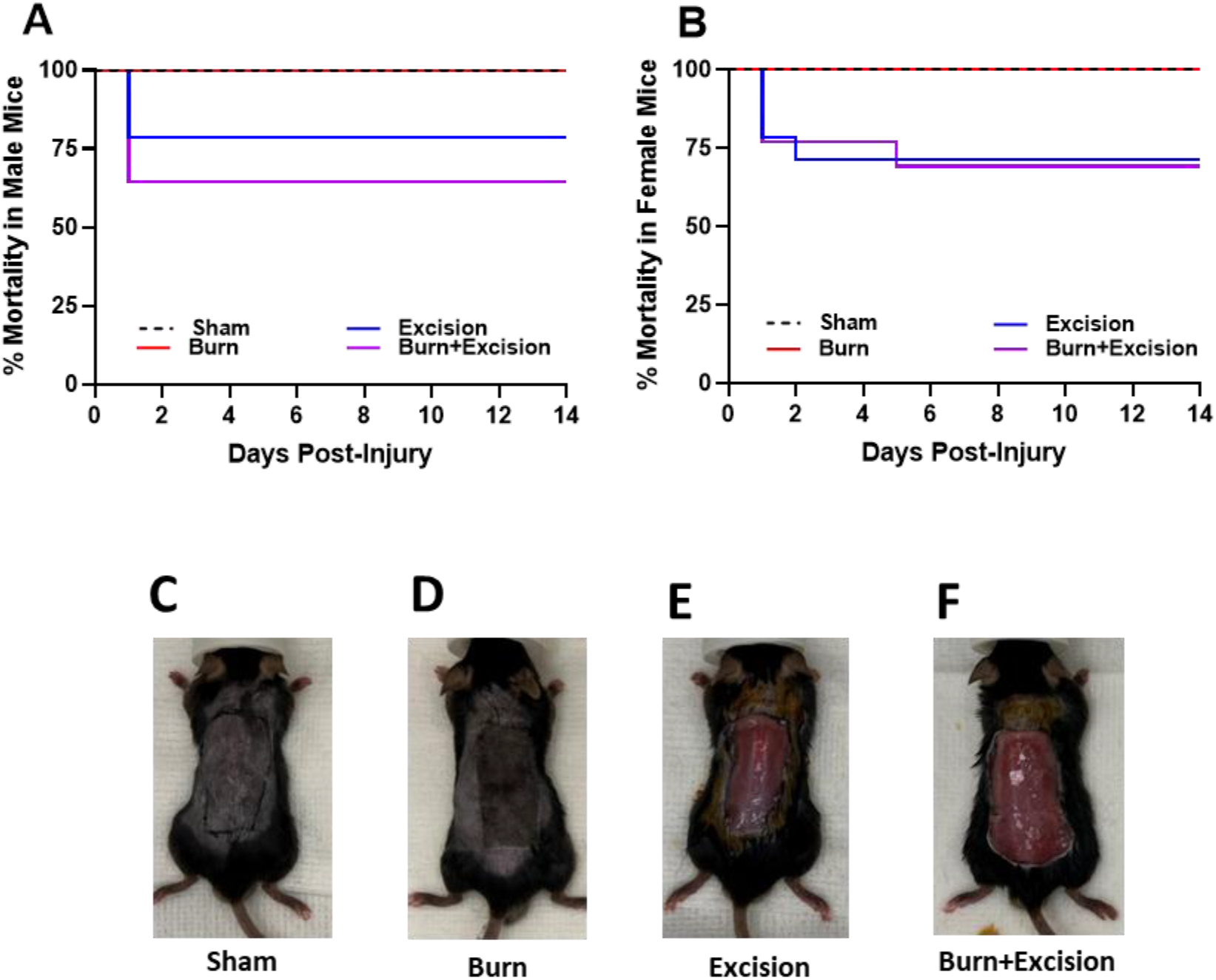
Mortality rate in male (A) and female mice (B). Data presented as the percentage of mice in each group that did not survive post procedure. Representative images of a mouse in the sham (C), burn only (D), excision only (E) and burn+excision (F) groups. n=10 in sham/burn only per sex. n=14 in excision only/ burn+excision per sex.

### Impact of Burn, Excision, and Combined Burn+Excision on Total Energy Expenditure and its Components

**Figure 3** shows a time course of absolute rates of REE, TEE and VH_2_O (Panels A-F) including both baseline and post injury periods. Panels G-L show percent change data from baseline for the same parameters plotted in (Panels A-F). Resting energy expenditure (REE) increased above baseline in all injury groups in both male (Time P<0.0001, Treatment P<0.0001, Two-Way ANOVA) (**Fig 3A**) and female mice (Time P<0.0001, Treatment P<0.0001, Two-Way ANOVA) (**Fig 3B**). Accordingly, REE was greater in all injury groups compared to sham mice post injury (Time P<0.0001, Treatment P<0.0001, Two-Way ANOVA) (**Figs 3A, B**). Irrespective of sex, the magnitude of the increase in REE post injury was greater in mice with excision and burn+excision compared to burn (Time P<0.0001, Treatment P<0.0001, Two-Way ANOVA) (**Figs 3A, B**). The magnitude and persistence of the increase in REE in the excision and burn+excision groups were comparable in both sexes. While there was an immediate increase in REE in all three injury groups, there was a more profound elevation in REE in excision and burn+excision groups (Time P<0.0001, Treatment P<0.0001, Two-Way ANOVA) (**Figs 3G, H**) that began to resolve at approximately 7-days post injury. In contrast, the more modest increase in post-injury REE in the burn group remained largely stable across the 14-day recovery period (**Figs 3G, H**). These dynamic changes in REE in response to different injury types were comparable in male and female mice.

**Figure 3.**
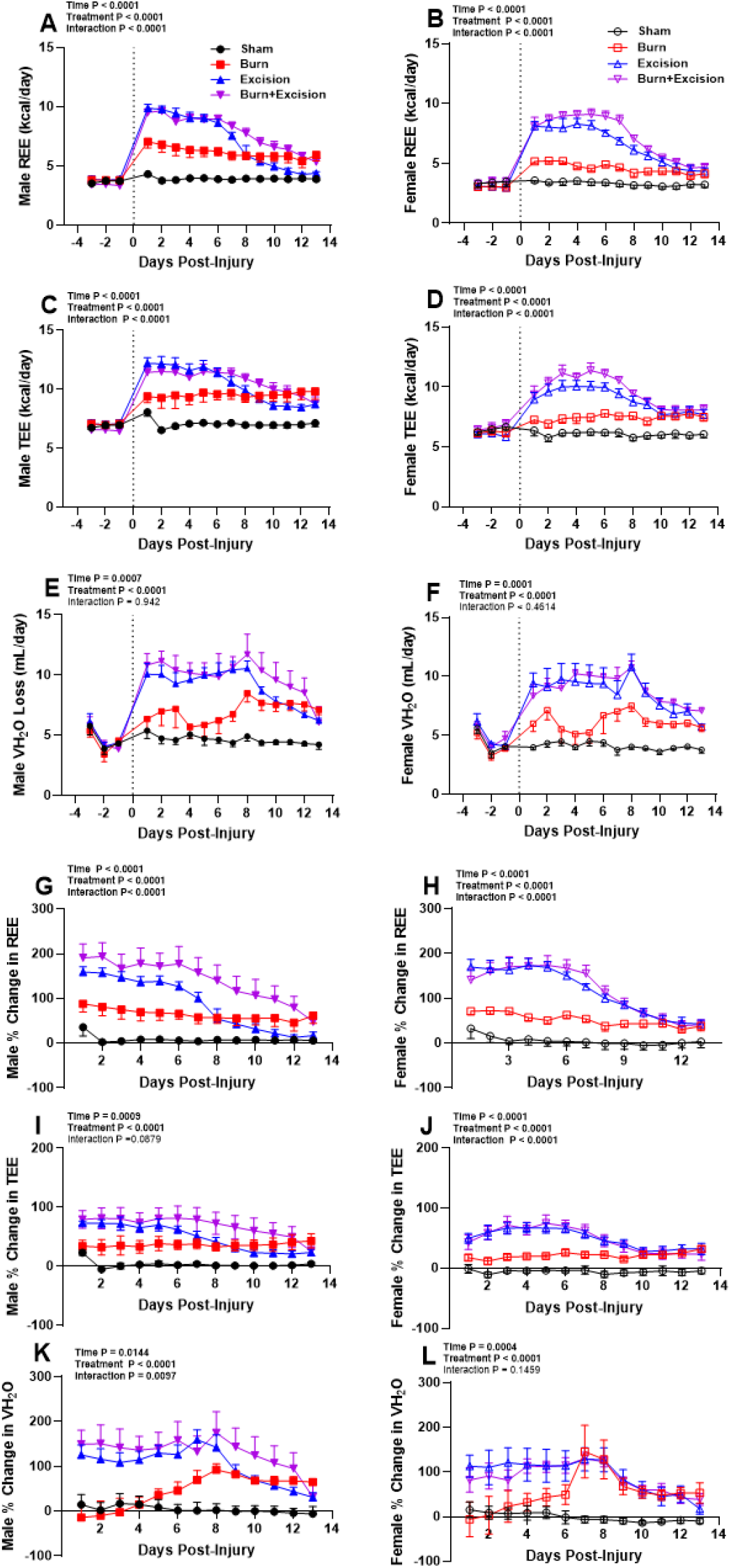
Resting energy expenditure (REE), total energy expenditure (TEE) and water vapor production rates (VH_2_0) in male (panels A, C and E) and female (panels B, D and F) mice. Data show three consecutive days of baseline measurements for all groups prior to procedures and 13 days of consecutive data starting at 6 p.m. on the day of procedures until 6 a.m. on the morning of euthanasia. Percent changes from baseline for REE, TEE, and water vapor production rates in male (panels G, I, and K) and female (panels H, J, and L) mice. Panels A-F show Two-Way ANOVA time, treatment and interaction P-values (excluding baseline data prior to injury). Panels G-L show percent change from baseline, analyzed by Two-Way ANOVAs with Tukey’s test for Multiple Comparisons. Values are group means±SEM, n=6-8 mice per group. REE is the lowest 30 min of EE measured in the light period and multiplied by 24 to express as kcal/day. REE was calculated from VO_2_ and VCO_2_ using the Weir equation.

Total energy expenditure (TEE) in male (**Fig 3C**) and female (**Fig 3D**) mice was elevated in all injury groups (Time P<0.0001, Treatment P < 0.0001). As with REE, the magnitude of the increase in TEE was greater in mice with excision and burn+excision compared to burn alone (Time P<0.0001, Treatment P<0.0001, Two-Way ANOVA) (**Figs 3I, J**). The magnitude and persistence of the increase in TEE in the excision and burn+excision groups were comparable in both sexes. Similar to TEE, marked increases in TEE in the excision and burn+excision groups began to dissipate at around 7-days post-injury, whereas the more modest increase in TEE in burn only mice remained quantitatively similar over the entire 14-days recovery period (Time P<0.0001, Treatment P<0.0001, Two-Way ANOVA) (**Figs 3I, J**).

Water vapor production rates in male (**Fig 3E**) and female (**Fig 3F)** mice were increased in all injury groups (Time P=0.0007, Treatment P<0.0001, Two-Way ANOVA). As with REE and TEE, the magnitude of this response was greater in mice with excision and burn+excision compared to burn only. Similarly, while the more modest increase in water vapor production rate was quantitatively stable across the 14-day recovery period in the burn group, a more pronounced elevation in water vapor production rate was observed in mice with excision and burn+excision that began to decline after the first 7-days post-injury (Time P≤0.0144, Treatment P<0.0001, Two-Way ANOVA) (**Figs 3K, L**).

**Figure 4** reports the absolute (Panels A-D) and relative (Panels E-H) contributions of REE to TEE in both the first and second weeks post injury. During the first week post-injury, REE was significantly greater in both sexes in the burn, excision, and burn+excision groups compared to sham (P<0.05, One-Way ANOVA) (**Fig. 4A and 4B**). In both sexes, REE in the excision and burn+excision group were also elevated compared to the burn groups in the first week post injury (P<0.01, One-Way ANOVA) (**Fig. 4A and 4B**). In female mice, REE in the burn+excision group remained elevated compared to the REE of the excision group in the first week post injury (P<0.01, One-Way ANOVA) (**Fig. 4B**). During the second week, the REE in male mice in the burn and burn+excision groups are elevated compared to sham mice (P<0.05, One-Way ANOVA) (**Fig. 4C**). The burn group also has elevated REE compared to the excision group in male mice during the second week post-injury (P<0.01, One-Way ANOVA) (**Fig. 4C**). In female mice, REE in the excision and burn+excision remained elevated compared to sham during the second week (P<0.05, One-Way ANOVA) (**Fig. 4D**). REE represented a significant proportion of the TEE in all injury groups compared to sham mice during the first week post-injury in male mice (P<0.0001, One-Way ANOVA) (**Fig. 4E**). In male mice, REE represented a greater proportion of TEE in the excision and burn+excision groups compared to burn alone (P<0.01, One-Way ANOVA) (**Fig. 4E**). In female mice, REE represented a greater proportion of TEE in the excision and burn+excision groups (but not the burn only group) compared to sham mice (P<0.001), though the burn+excision group was also significantly different to the burn alone group (P<0.001, One-Way ANOVA) (**Fig. 4F**). In the second week post injury, the relationship between REE and TEE was comparable between sham and injury groups in both sexes (**Fig. 4G and 4H**).

**Figure 4:**
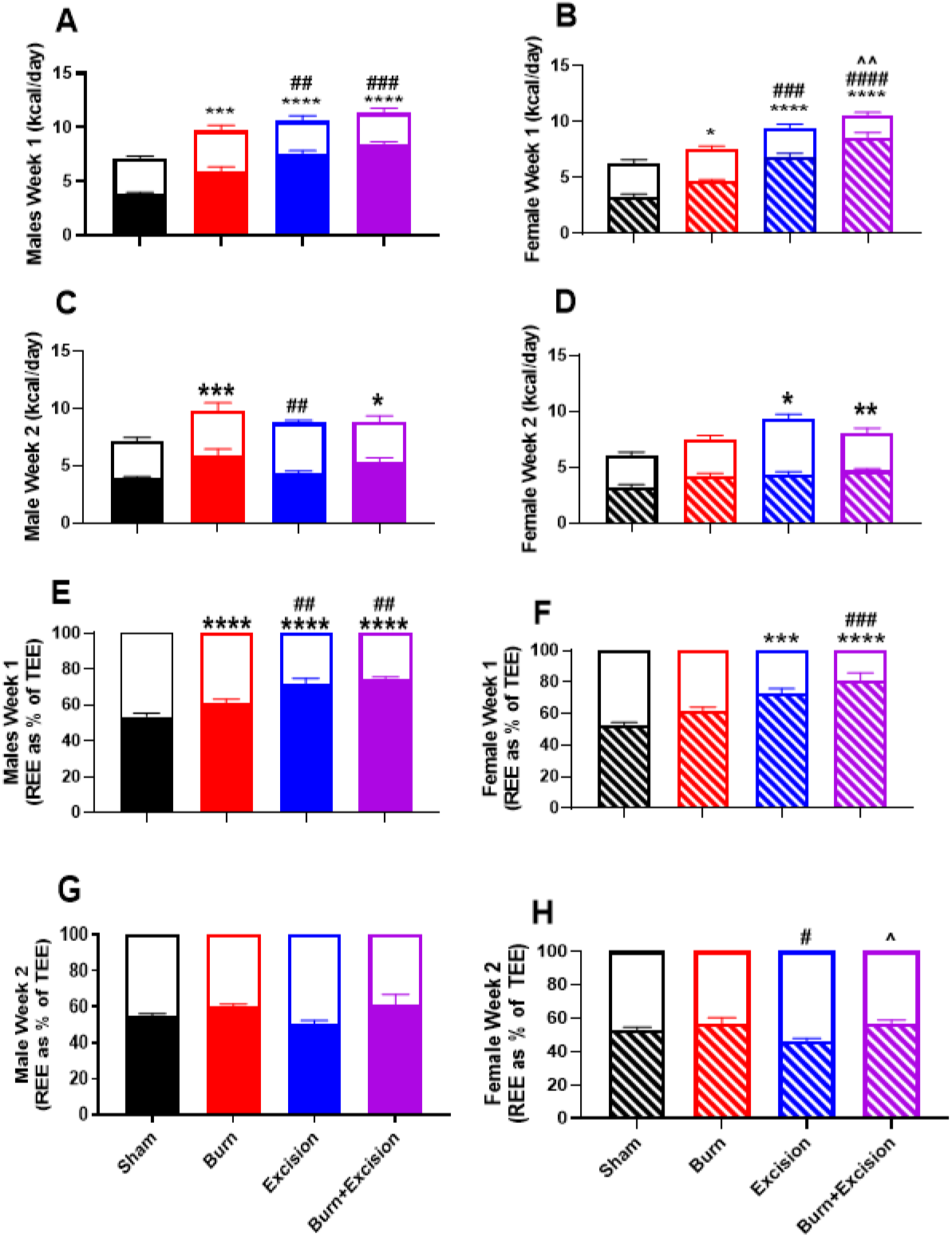
Average daily REE (colored bars) and TEE (unfilled bars) in male mice during the first (A) and second (C) week post-injury and female mice during the first (B) and second (D) week post-injury. All tests in A-D denotes differences in REE. Daily REE (colored bars) as a percentage of TEE (unfilled bars) in male mice during the first (E) and second (G) week post-injury and in female mice during the first (F) and second (H) week post-injury. Values are means ± standard errors, n=5-7 per group. All panels show REE analyzed using One-Way ANOVAS, with Tukey’s test for Multiple Comparisons. *P<0.05, **P<0.01, ***P<0.001 and ****P<0.0001 vs. sham REE. #P<0.05, ##P<0.01 and ###P<0.001 vs. burn only REE. ^^P<0.01 and ^^^P<0.001 vs. excision only REE.

### Impact of Burn, Excision, and Combined Burn+Excision on Behavior

In male mice, cumulative food intake was greater in all injury groups, but this was primarily observed in the second week post-injury (Treatment effect P=0.0009, Two-Way ANOVA) (**Fig. 5A**). When examining each week individually, male mice only demonstrated a significant difference in daily food intake in the second week post injury (**Fig. 5C**, P<0.05 One-Way ANOVA). In female mice, cumulative food intake was greater in mice with excision and burn+excision injuries (Treatment effect P<0.0001, Two-Way ANOVA), where female mice in the excision and burn+excision groups ate more food than sham during the 2-week recovery period. As with male mice, this was driven by hyperphagia in the second week of recovery (**Fig. 5F**, P<0.05 One-Way ANOVA). With respect to water intake, male burn+excision mice exhibited a significant treatment effect (Treatment effect P<0.0001, Two-Way ANOVA) (**Fig. 5G**). While there is a significant increase in average daily water intake in the male excision and burn+excision mice during the first week post-injury (P<0.05, One-Way ANOVA) (**Fig 5H**), all groups had comparable water intake by week two (**Fig. 5I**). In female mice, there was a significant treatment effect on cumulative water intake (Treatment P<0.0001, Two-Way ANOVA), where cumulative water intake was greater in all three injury groups ( **Fig. 5J**). Indeed, in both the first and second week post-injury, female excision and burn+excision groups had significantly greater average daily water intake compared to sham (P<0.05, One-Way ANOVA) (**Fig. 5J and 5L**). Additionally, during the second week, burn alone is also shown to have increased water intake in female mice compared to the sham group (P<0.05, One-Way ANOVA) (**Fig. 5L**).

**Figure 5:**
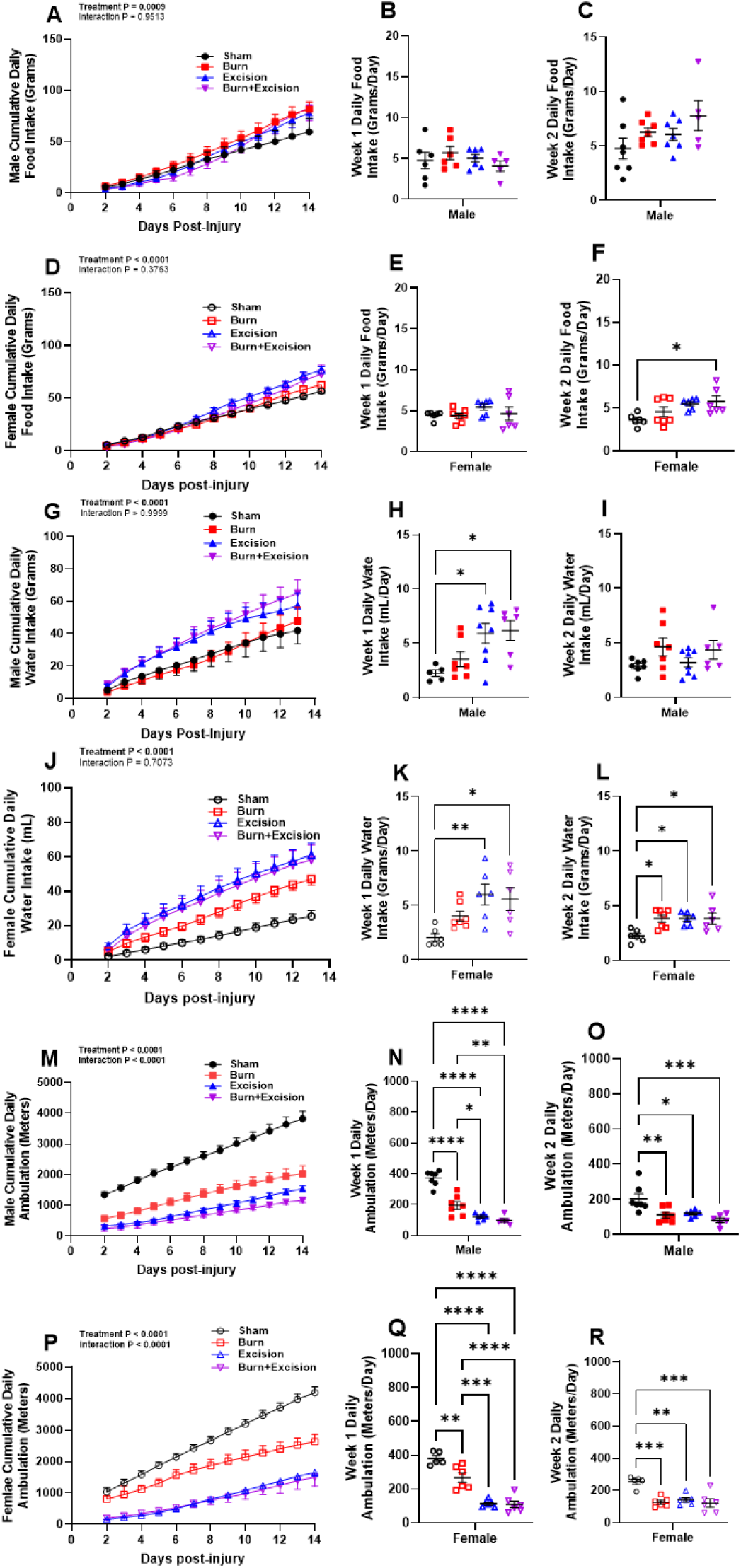
Cumulative food intake in male (A) and female (D) mice. Average daily food intake for male mice during the first (B) and second (C) week post-injury, and in female mice during the first (E) and second (F) week post-injury. Cumulative water intake in male (G) and female (J) mice. Average daily water intake for male mice during the first (H) and second (I) week post-injury, and in female mice during the first (K) and second ( L) week post-injury. Cumulative ambulation in male (M) and female (P) mice. Average daily ambulation for male mice during the first (N) and second (O) week post-injury, and in female mice during the first (Q) and second (R) week post-injury. Values are means±SEM, n=6-8 per group. Panels A, D, G, J, M, and P analyzed using a Two-Way ANOVA. A One-Way ANOVA was used to analyze data in all other panels, with Tukey’s test for Multiple Comparisons were used to compare groups. *P<0.05, **P<0.01, ***P<0.001 and ****P<0.0001 vs. sham.

In male and female mice, cumulative cage ambulation was markedly lower in all injury groups (Treatment effect P<0.0001, Two-Way ANOVA) (**Fig. 5M and 5P**). Irrespective of sex, average daily ambulation in both week 1 (**Fig. 5N and Q**) and week 2 (**Fig. 5O and 5R**) post-injury was lower in all injury groups (P<0.01, One-Way ANOVA), where this injury effect was greatest in the excision and burn+excision groups. During the first week post-injury, the excision and burn+excision groups ambulation is significantly lower compared to burn alone (P<0.01, One-Way ANOVA) (**Fig. 5N and Q**). In female mice, starting at 5 days post-injury, the cumulative movement for the burn mice is significantly lower than the sham mice ( **Fig. 5P**).

### Impact of Burn, Excision and Combined Burn+Excision on Body Mass and Composition

Percent change in body mass is shown in **Fig. 6**. Body mass (corrected for the mass removed by wound excision) significantly decreased in the first two days post-injury in the excision and burn+excision groups, regardless of sex (Time P<0.0001, Treatment P≤ 0.0002, Two-Way ANOVA) (**Fig. 6A and B** (raw data are reported in **Supplemental Fig. 3A and 3B**)). Both male and female mice in the excision and burn+excision groups had a greater loss in body mass compared to the burn group at 2-days post injury (P<0.05) (**Fig. 6C and D** (raw data are reported in **Supplemental Fig. 3C and D**)). In female mice, the excision group also had lower body masses compared to sham mice at 2-days post injury (P<0.05, One-Way ANOVA) (**Fig 6D**). In male mice, the burn+excision group had significantly lower body masses compared to both the sham and burn groups at 2-days post injury (P<0.05, One-Way ANOVA) (**Fig 6C**). While the acute change in body mass associated with excision and burn+excision was sustained in male mice at both 7-(P<0.01, One-Way ANOVA) and 14-days post injury (P<0.001, One-Way ANOVA) (**Fig. 6E and 6G**), female mice in the excision and burn+excision groups were comparable to sham and burn mice by day 7 post injury (**Fig. 6F and 6H**).

**Figure 6:**
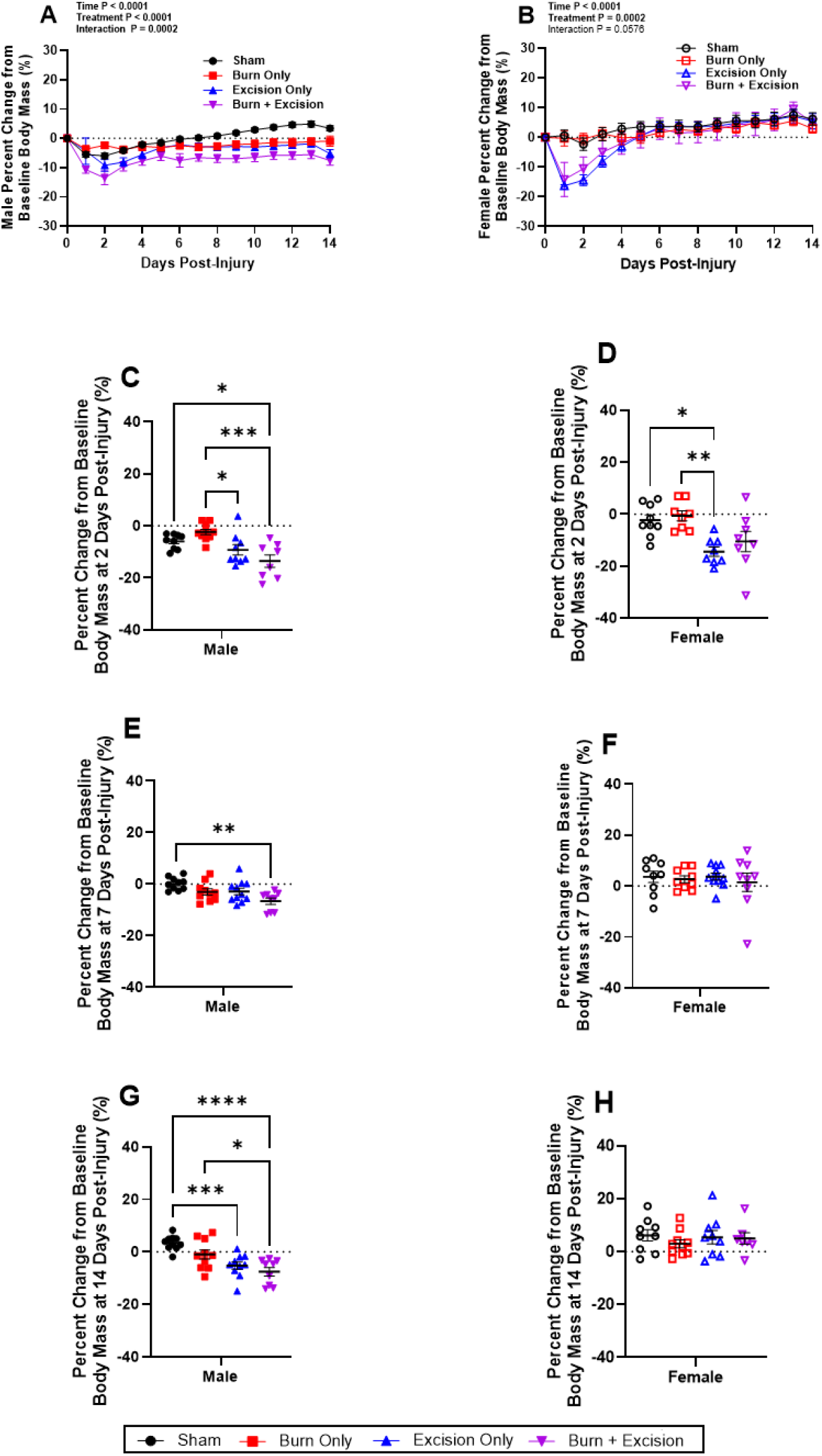
Percent change from baseline body mass in male (A) and female (B) mice. Data show percent change in body mass from day 0 prior to procedure and during the 14-day recovery period. Percent change in body mass is also shown at days 2, 7, and 14 post-injury in male (panels C, E, and G) and female (panels D, F, and H) mice. Values are means±SEM, n=9-11 per group. Panels A and B analyzed by Two-Way ANOVA, with Tukey’s test for Multiple Comparisons comparing between groups over time. On all other panels, One-Way ANOVA, with Tukey’s test for Multiple Comparisons were used to compare groups. *P<0.05, **P<0.01, ***P<0.001 and ****P < 0.0001.

Change in body composition is shown in **Fig. 7**. Male mice with excision or burn+excision injuries had reduced lean mass compared to sham mice at the end of the 14-day recovery period (P<0.01, One-Way ANOVA) (**Fig. 7A** (raw data are reported in **Supplemental Fig. 4A**)). Similarly, absolute fat mass was lower in male excision and burn+excision groups relative to sham, and was also lower in the burn group compared to the sham group (P<0.01, One-Way ANOVA) (**Fig. 7B** (raw data are reported in **Supplemental Fig. 4B**). Female mice showed no significant change in lean or fat mass compared to pre-injury in any group (**Fig. 7B and 7D** (raw data are reported in **Supplemental Fig. 4D**). Regardless of sex, the excision and burn+excision groups had lower bone mineral density compared to sham (P<0.01, One-Way ANOVA), whereas bone mineral density was not lower in burn alone versus sham groups (**Fig. 7E and 7F, Supplemental Fig. 4E and 4F**). In addition, compared to burn alone, bone mineral density was lower in the burn+excision group in male mice and the excision group in female mice (P<0.01, One-Way ANOVA) (**Fig. 7E and 7F**). Bone mineral content was also lower in the male burn+excision group compared to sham, burn, and excision groups (P<0.05, One-Way ANOVA) (**Fig 7G**). In female mice, both the excision and burn+excision groups had significantly lower bone mineral content (P<0.01, One-Way ANOVA) (**Fig 7H**).

**Figure 7:**
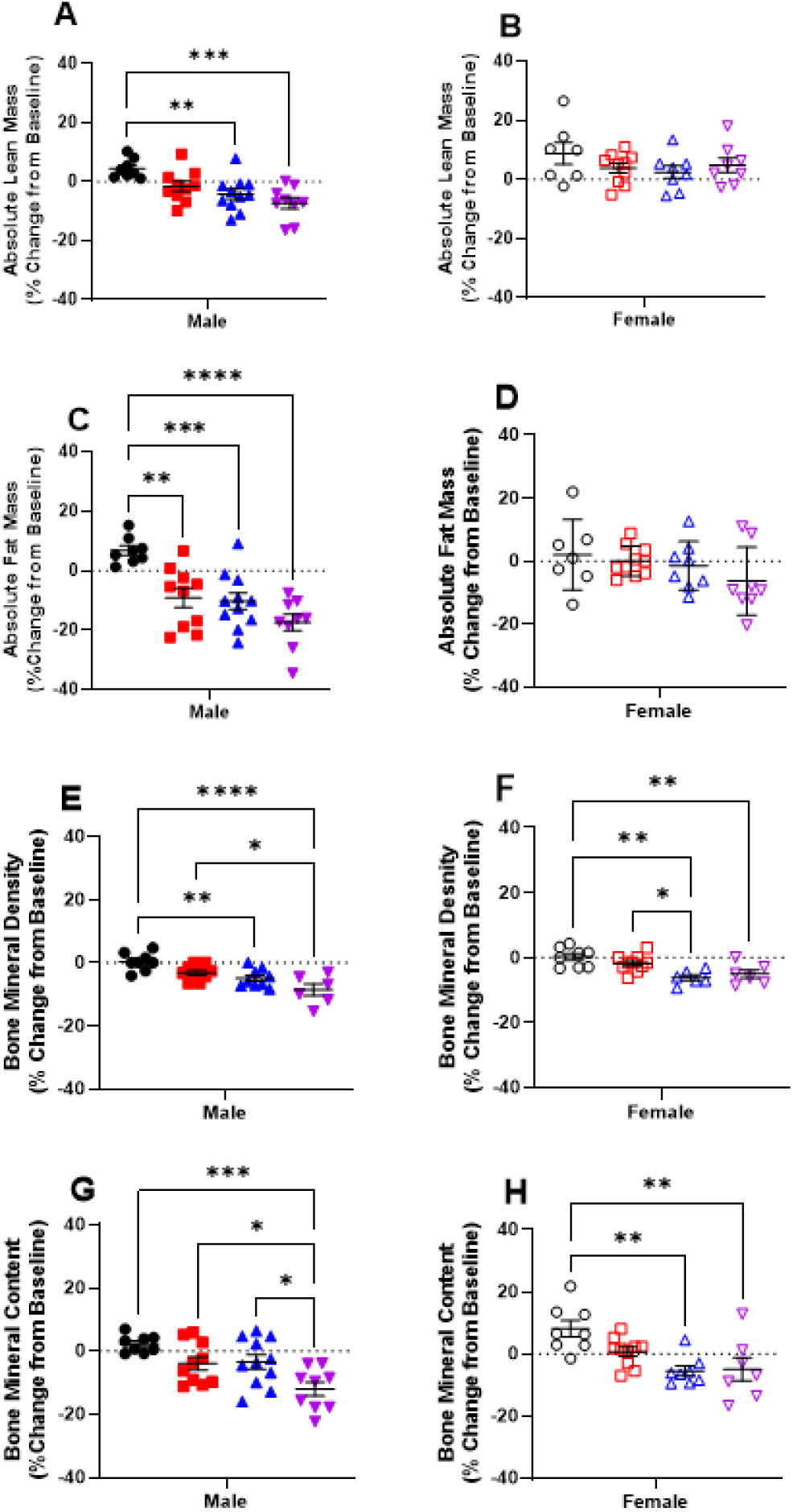
Percent change from baseline lean mass in male (A) and female (B) mice. Percent change from baseline fat mass in male (C) and female (D) mice. Percent change from baseline bone mineral density in male (E) and female (F) mice. Percent change from baseline bone mineral content in male (G) and female (H) mice. Data show percent change in masses compared from day 0 prior to procedure compared to the morning of euthanasia obtained from DEXA scans. Values are means±SEM, n=9-11 per group. One-Way ANOVA, with Tukey’s test for Multiple Comparisons were used to compare groups. *P<0.05, **P<0.01, ***P<0.001 and **** P<0.0001.

### Impact of Burn, Excision, and Combined Burn+Excision on Organ Mass

Tissues and organ masses were normalized to excision-corrected body mass at the time of euthanasia. In male mice, the relative liver size was significantly higher in the burn+excision group compared to all other groups (P<0.05, One-Way ANOVA) (**Fig. 8A**, (raw data are reported in **Supplemental Fig. 5A**)). Relative liver size was greater in all injury groups compared to sham in female mice (P<0.05, One-Way ANOVA) (**Fig 8B**). In female mice, liver mass was elevated in the burn+excision group compared to both the sham and burn alone groups (P<0.01, One-Way ANOVA) (**Fig. 8B** (raw data are reported in **Supplemental Fig. 5B**)). In both sexes, burn+excision mice had higher relative heart masses compared to their sham counterparts (P<0.05, One-Way ANOVA) (**Fig. 8C and 8D** (raw data are reported in **Supplemental Fig. 5C and 5D**)). In male mice, the excision group exhibited greater relative heart mass compared to sham mice (P<0.01, One-Way ANOVA) (**Fig. 8C**). Excision and burn+excision groups had a higher relative kidney mass compared to both their sham and burn only counterparts in male mice (P<0.05, One-Way ANOVA) (**Fig. 8E** (raw data are reported in **Supplemental Fig. 5E**)). In female mice, relative kidney mass was elevated in the burn+excision group compared to the sham and burn alone groups, while the excision alone group only showed greater relative kidney mass when compared to sham (P<0.05, One-Way ANOVA) (**Fig. 8F** (raw data are reported in **Supplemental Fig. 5F**)). When comparing relative spleen masses, burn+excision groups had significantly greater spleen masses compared to all other groups, regardless of sex (P<0.05, One-Way ANOVA) (**Fig. 8G and 8H** (raw data are reported in **Supplemental Fig. 8G and 8H**). In male mice, excision only showed a greater relative spleen mass than both sham and burn only mice (P<0.05, One-Way ANOVA) (**Fig. 8G**). Burn alone male mice also had greater spleen mass compared to sham (P<0.01, One-Way ANOVA) (**Fig. 8G**). In female mice, excision only spleen mass was significantly greater when compared to the sham mice (P<0.05, One-Way ANOVA) (**Fig. 8H**).

**Figure 8:**
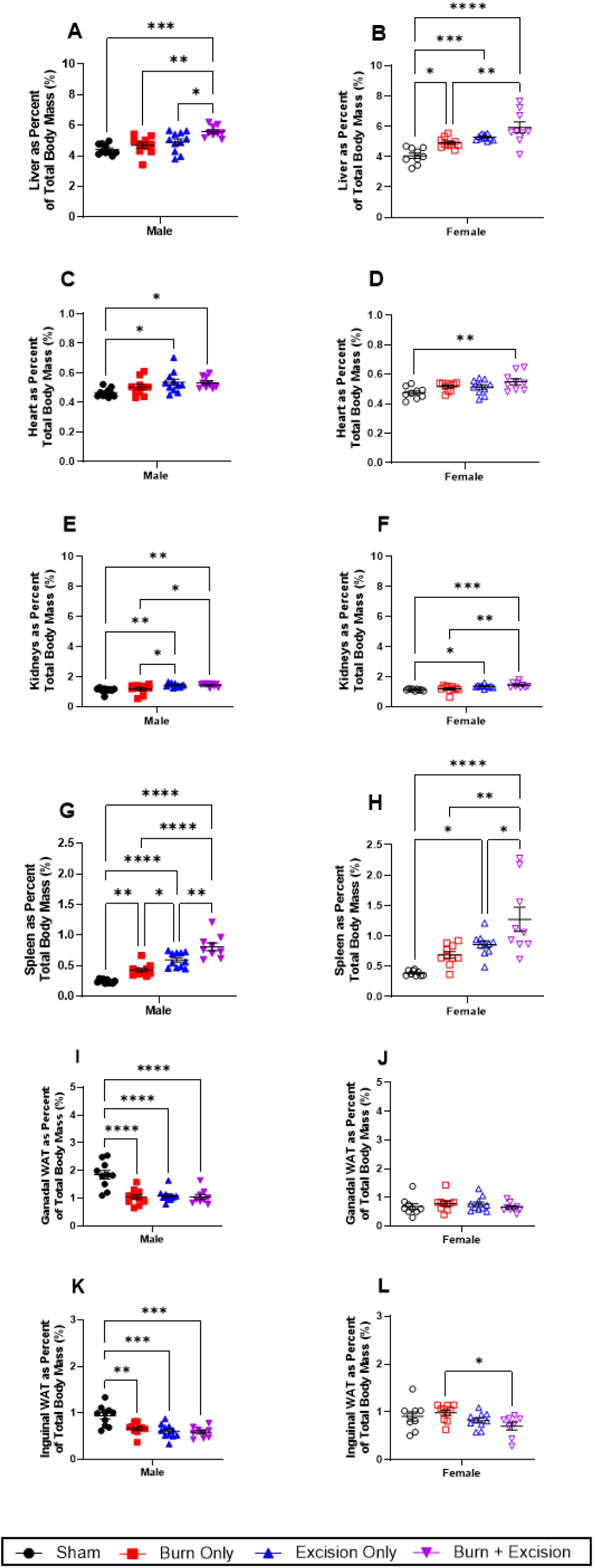
The relative liver size in male (A) and female (B) mice. The relative heart size in male (C) and female (D) mice. The relative kidneys size in male (E) and female (F) mice. The relative spleen size in male (G) and female (H) mice. The relative gonadal white adipose tissue depot size in male (I) and female (J) mice. The relative inguinal white adipose tissue depot size in male (K) and female (L) mice. All organs taken at day 14 immediately after euthanasia. Values are means ± standard errors, n = 9-11 per group. One-Way ANOVAS, with Tukey’s test for Multiple Comparisons were used to compare groups. * = P<0.05, ** = P<0.01, ***= P<0.001 and **** = P < 0.0001.

In line with our DEXA data, male mice had lower relative gondal adipose tissue depot mass in all injury groups (P<0.0001, One-Way ANOVA) (**Fig. 8I** (raw data are reported in **Supplemental Fig 5I**)). Similarly, male mice had smaller inguinal white adipose tissue depots in all injury groups compared to sham mice (P<0.01, One-Way ANOVA) (**Fig. 8K** (raw data are reported in **Supplemental Fig 5K**)). Female mice do not show significant differences between relative gonadal white adipose tissue depot mass (**Fig. 8J** (raw data are reported in **Supplemental Fig 5J**)), In female mice, the inguinal white adipose tissue depot was significantly smaller in the burn+excision group compared to burn alone group (P<0.05, One-Way ANOVA) (**Fig 8L** (raw data are reported in **Supplemental Fig 5L**)).

### Impact of Burn, Excision, and Combined Burn+Excision on Blood Cell Composition

Following euthanasia and blood collection, a complete blood count with differential was carried out. In male mice, all injury groups demonstrated lowered red blood cell count compared to sham (P<0.01, One-Way ANOVA) (**Fig. 9A**), where male burn+excision exhibited a lower red blood cell count compared to burn alone (P<0.05, One-Way ANOVA) (**Fig. 9A**). In female mice, only the burn+excision group had significantly lower red blood cell counts compared to sham (P<0.05, One-Way ANOVA) (**Fig. 9B**). Similarly, male hematocrit and hemoglobin levels were lower in all injury groups compared to sham (P<0.01, One-Way ANOVA) (**Fig. 9C and 9E**). In both male and female mice, hematocrit and hemoglobin were lower in the burn+excision group compared to burn alone (P<0.01, P<0.05, One-Way ANOVA) (**Fig. 9C-9F**). In both sexes, excision and burn+excision groups had elevated platelet counts compared to sham (P<0.05, One-Way ANOVA) (**Fig. 9G and 9H**). Male mice in the burn group also had an elevated platelet count compared to sham (P<0.01, One-Way ANOVA) (**Fig. 9H**). Regardless of sex, the mean platelet volume was greater in the burn+excision groups compared to sham (P<0.01, One-Way ANOVA) (**Fig. 9I and 9J**).

**Figure 9:**
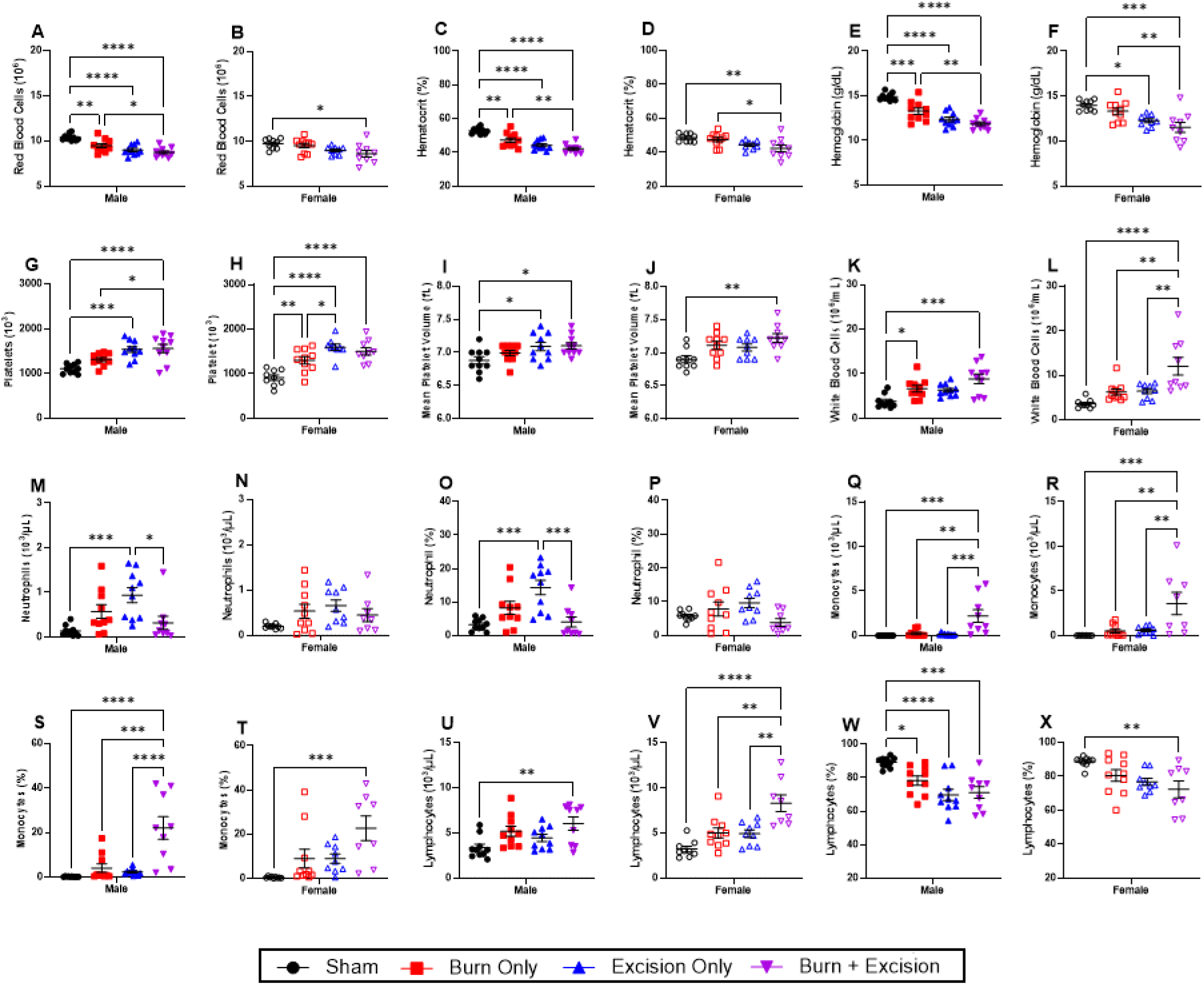
Red blood cell count in male (A) and female (B) mice. Hematocrit in male (C) and female (D) mice. Hemoglobin concentration in male (E) and female (F) mice. Platelet count in male (G) and female (H) mice. Mean platelet volume in male (I) and female (J) mice. Total white blood cell count in male (K) and female (L) mice. Neutrophil number in male (M) and female (N) mice. Percent of neutrophils in male (O) and female (P) mice. Monocyte number in male (Q) and female (R) mice. Percent of monocytes in male (S) and female (T) mice. Lymphocyte number in male (U) and female (V) mice. Percent of lymphocytes in male (W) and female (X) mice. All blood samples were collected on day 14 post injury immediately after euthanasia. Values are means ± standard errors, n=9-11 per group. One-Way ANOVAS, with Tukey’s test for Multiple Comparisons were used to compare groups. *P<0.05, **P<0.01, ***P<0.001 and ****P<0.0001.

Male mice had elevated white blood cell counts in the burn alone and burn+excision groups compared to their sham counterparts (P<0.05, One-Way ANOVA) (**Fig. 9K**). In female mice, the burn+excision group had elevated white blood cell counts compared to all other groups ( P<0.01, One-Way ANOVA) (**Fig. 9L**). In male mice, the excision alone gourp had elevated absolute and relative neutrophil levels compared to both sham and burn+excision mice (P<0.05, One-Way ANOVA) (**Fig. 9M and 9O**). Neither the absolute neutrophil count nor the ratio of neutrophils in the blood was significantly changed in female mice ( **Fig. 9N and 9P**). In both sexes, absolute monocyte count elevated in the burn+excision group compared to all other groups (P<0.01, One-Way ANOVA) (**Fig. 9Q and 9R**). Relative monocyte count was also greater in the male and female burn+excision groups compared to sham (P<0.001, One-Way ANOVA) (**Fig. 9S and 9T**). Relative monocyte counts were also significantly elevated in the burn+excision group compared to burn and excision alone in male mice only (P<0.001, One-Way ANOVA) (**Fig. 9S**). Absolute lymphocyte counts were elevated in the burn+excision groups, regardless of sex (P<0.01, One-Way ANOVA) (**Fig. 9U and 9V**). Female burn+excision absolute lymphocyte counts were also elevated compared to the burn and excision alone groups ( P<0.01, One-Way ANOVA) (**Fig 9V**). Relative lymphocyte abundance was lower in the burn+excision compared to sham, irrespective of sex (P<0.01, One-Way ANOVA) (**Fig. 9W and 9X**).

## Discussion

Patients with severe burns endure a prolonged hypermetabolic stress response that can be accompanied by profound cachexia. Data suggest that this hypermetabolic response peaks in the first 2-3 weeks following burn injury^18–21^. Further, additional clinical data suggest that burn wound excision itself may be a driver of burn-induced hypermetabolism in burn survivors^8^. Despite this, mouse models of burn injury typically do not incorporate burn wound excision, which may limit the ability of these models to accurately recapitulate the hypermetabolic-catabolic stress response to burns observed in patients. In this study, we assessed the effect of scald burn and wound excision alone and in combination on hypermetabolism and cachexia in male and female mice acclimated to thermoneutral housing. By performing body composition scans and comprehensive metabolic phenotyping following sham, burn, wound excision, and burn followed by excision injuries, we found that wound excision alone results in profound hypermetabolism and cachexia, exceeding that seen in burn injury alone and broadly comparable to that seen in animals with combined burns and wound excision. Indeed, the acute hypermetabolic response observed in mice that underwent excision alone was comparable to mice that underwent burn+excision. However, burn injury served to exacerbate the persistence of this hypermetabolic response and the degree of cachexia when comparing the excision and burn+excision groups. These data support observations in burn patients, suggesting burn wound excision contributes to acute hypermetabolism, thereby underscoring the importance of wound excision when attempting to accurately model burn-induced hypermetabolism and catabolism in mice.

We recently reported the lack of standardized reporting on mouse total body surface area (TBSA) and absolute burn size in burn rodent literature, where most studies report relative burn size alone^22^. This makes reproducibility a challenge and likely means that the relative burn size reported in some studies may be inaccurate. After establishing a new refined approach to determine TBSA in mice that accounts for the presence of the *panniculus carnosus*, which is largely absent in humans, we developed revised Meeh constants for male and female mice that allow TBSA estimation from body mass^15^. In doing so, we estimate that the ∼27g male and ∼19g female mice used in the current study had TBSA of ∼60cm^2^ and ∼45cm^2^, respectively. By creating wounds that were ∼8.6cm^2^ (males) and ∼6.5cm^2^ (females), this computed to full thickness wound sizes that involved approximately 13-15% of the TBSA. Burn only animals exhibited modest hypermetabolism and no mortality when subjected to ∼15% TBSA full thickness burn injuries. Iin mice that underwent wound excision alone or burn injury with subsequent wound excision, we observed approximately 25-30% mortality. These deaths typically occurred within the first 24-48 hours post-injury, despite wound excisions resulting in little to no bleeding. Upon necropsy, we were unable to find a clear explanation (e.g. organ damage) for the loss of animals that underwent wound excision. That said, these data suggest that the inclusion of wound excision in a mouse model of scald burns markedly elevates the severity of this model. We posit that, in mice, a burn encompassing 15% of the TBSA (that is subsequently excised) reflects a model of severe burn trauma.

We observed an acute hypermetabolic response in all injury groups evidenced by elevated resting energy expenditure (REE); however, excision alone and burn+excision resulted in a more severe acute hypermetabolic response when compared to both sham and burn alone, regardless of sex. Indeed, REE was double that of the sham group in both excision and burn+excision groups in the first week post burn, irrespective of sex. Relating this to each group’s baseline data, REE was ∼48% above normal in the burn groups, ∼100% above normal in the excision groups, and ∼135% above normal in the burn+excision groups. These data show that while burn alone resulted in hypermetabolism, wound excision markedly acerbates the hypermetabolic response to burns. We have previously reported a modest hypermetabolic response to burns by employing continuous REE measurements post-injury in mice with scald burns (where wound excision was not performed) ^23^. Our current REE data quantitatively differ from our previous data ^23^ in terms of the magnitude of this response. In our previous report, mice were housed at 24°C prior to experimentation, then randomized to either standard (24°C) or thermoneutral (30°C) vivarium temperatures during their recovery from sham/burn procedures. Modest hypermetabolism was only apparent in mice housed at thermoneutrality, suggesting existing cold stress due to sub-thermoneutral housing masks burn-induced hypermetabolism in mice^23^. In our current study, mice were acclimated to thermoneutrality for 4 weeks prior to burn injury alone exhibited a more profound hypermetabolic response than those in our prior study, where mice were not acclimated to thermoneutrality before burn injury ^23^. This suggests acute housing at 30°C post burn injury may be insufficient to adequately model burn-induced hypermetabolism in mice. Rather, long-term acclimation to thermoneutral housing to fully humanize mouse energetics is necessary to more appropriately study the stress response to burns.

The profound hypermetabolic response observed in the excision and burn+excision groups translated to greater TEE in these groups relative to both the sham and burn alone groups, particularly in the first week post-injury. In all injury groups, greater REE drove increased TEE, where in relative terms, REE represented a greater proportion of TEE, particularly in the f irst week post burn. This corroborates human data where respiratory gas exchange and daily measures of whole-body carbon dioxide production indicate the hypermetabolic response to burns results in elevations in both REE and TEE, as well as the relative contribution of REE to TEE^24^. This spike in REE persists for the 14 days of the study, though it begins to taper off approximately one-week post-injury. While all injury groups remained hypermetabolic throughout the 14-day recovery period, the magnitude of the increase in REE and TEE was largely stable in animals with burn injuries alone. The profound increase in REE (and thus TEE) seen in animals with excision alone and burn+excision remained stable for the first 7-8 days post-injury, before beginning to decline towards that seen in the burn only group.

It is known that acute hypermetabolism in burn patients is a function of evaporative and convective heat loss from wounds^10^ where there is a strong positive relationship between area burned and water loss^9^. This indicates heat loss from burn wounds is a key driver of burn-induced hypermetabolism. Further, excision of burn wounds themselves, even if delayed for several weeks from the initial injury, triggers profound hypermetabolism in burn patients^8^. In our study, we continuously measured the rate of whole-body water loss through respiratory gas exchange, noting the profound impact of wound excision ( in either the absence or presence of burn injuries) on body water loss. Indeed, traces of daily water loss for all injury groups were remarkably similar to traces of daily REE and TEE, again indicating that heat loss is a significant driver of burn-induced hypermetabolism. Within the three injury groups, increased body water loss was more modest in the burn only group, where we posit that the presence of an eschar minimizes wound water loss in mouse models of severe burns. Indeed, we noted a spike in body water loss at ∼6-8 days post injury in both male and female mice with burn injuries alone, which corresponds to when eschars begin to separate and slough off burned mice. This supports the notion that the presence of the eschar in mice with burn injuries alone limits acute water loss and thus hypermetabolism post injury.

In addition to allowing high-resolution monitoring of respiratory gas exchange and thus determination of REE, TEE, and body water turnover, contemporary metabolic phenotyping cages also support the automated measurement of food and water consumption, as well as the animals’ movement around their cage in real time. While a concern in rodent models of burn trauma is that acute cachexia may be compounded by suppressed food intake post-injury, our data suggests that in both males and female, food intake was comparable across all groups in the first week post-injury. Moreover, there was a significant treatment effect of injury on food intake in both males and females when comparing daily food intake in the second week post-injury, where there was a hyperphagic response in excision and burn+excision groups, suggesting increased energy intake in response to greater TEE. In line with marked increases in whole-body water loss, there was a significant treatment effect on cumulative water intake, where male and female mice with excision and burn+excision consumed almost twice as much water than sham treated mice. As might be expected, there was a profound injury effect on cage ambulation. This was particularly apparent in both male and female randomized to excision and burn+excision, where in the first week post-injury, cage ambulation was ∼75% lower than the corresponding sham group. Burn injury alone also resulted in a marked reduction in cage ambulation compared to sham treated animals, this effect was less pronounced when compared to the excision and burn+excision groups. While the injury effect on cage ambulation persisted across the entire 14-day recovery period, as expected, it was more pronounced in the first week post-injury. Since ambulation contributes to TEE, this suggests that energy expenditure related to cage ambulation was lower in mice with injuries, particularly those in the excision and burn+excision groups. This, and the profound increase in REE in these groups, likely underlays the shift in the contribution of REE to TEE, where greater REE driven by heat loss and a heighted ATP demand for healing response occurs in the context of lower non-REE related energy expenditure (i.e., ambulation).

While the response to injury in terms of whole body energetics and related feeding and movement behaviors were broadly similar in male and female mice, we observed clear sexual dimorphism in terms of body mass and composition changes in response to injury. Both male and female mice in the excision and burn+excision groups underwent profound losses in body mass in the first 2-3 days post-injury (even when accounting for the loss of mass from wound excision). However, female mice in all three injury groups recovered their body mass during the 14-day recovery period. In contrast, cachexia persisted in male mice in the excision and burn+excision groups, where body mass remained significantly (∼10-15%) lower that baseline at 14-days post injury. This persistent cachexia in males in response to injury was also apparent when comparing baseline and 14-day post-injury body composition scans, where changes in absolute fat and lean mass were apparent in male mice with excision and burn+excision, but not in female mice. Indeed, this sexual dimorphism was also apparent at necropsy, where male mice in all injury groups had lower gonadal and inguinal fat pad masses (relative to total mass) compared to sham. Conversely, relative fat pad masses in female mice were comparable to sham across all three injury groups. Subtle differences in acute REE and feeding response to injury may underlay this apparent sexual dimorphism. Indeed, while the impact of injury on REE and TEE was broadly similar in males and females, the relative increase in both REE and TEE in response to excision and burn+excision were numerically greater in male compared to female mice in the first week post-injury. Further, cumulative food intake in females was greater in the excision and burn+excision groups relative to sham, whereas in males, while there was a hyperphagic response to excision and burn+excision in the second week post-injury, cumulative food intake over the 14-day recovery period was comparable to sham. When considering that female mice are significantly smaller than male mice, food intake per gram of body mass was ∼30% greater in female mice in the excision and burn+excision groups when comparing to the corresponding male groups. Collectively, greater relative hypermetabolism combined with a less pronounced increase in food intake (relative to body mass) contributes to a persistent energy deficit in male mice, explaining the sexual dimorphic response to injury in terms of cachexia.

While we observed clear sexual dimorphism in terms of lean and adipose tissue cachexia in response to injury, we observed similar levels of bone reabsorption in both male and female mice. Indeed, relative to sham, there was a marked loss of bone mineral content and density in both the excision and burn+excision groups, irrespective of sex. In line with our REE, TEE, and water loss data, this impact of excision or burn+excision on bone reabsorption was considerably more pronounced than that of burn injury alone. Beyond hypermetabolism and bone reabsorption, we also saw pronounce hepatomegaly and splenomegaly in excision and burn+excision groups, irrespective of sex. Greater absolute and relative liver and spleen masses were most pronounced in the burn+excision groups, even when compared to excision only groups, underscoring that excision of burn wounds increases the severity of burn models and that the combination of burns and subsequent excision may be additive with respect to organ dysfunction. Further underscoring the impact of wound excision on the severity of mouse models of burn injury, our complete blood count with white cell differential data at 14 days post-injury showed greater reductions in red blood cell number. Percent hematocrit and hemoglobin concentration were markedly lower in excision and burn+excision relative to both sham and burn only groups, irrespective of sex. Further, while there was an increase in white blood cell count in response to all injury groups, this response was most pronounced in mice with combined burn+excision injuries. Similarly, we observed a greater increase in absolute and relative monocyte concentration only in mice with combined burn+excision injuries, suggesting that immune response to burn+excision are more pronounced when compared to burn or excision injuries alone.

While inclusion of wound excision in mouse models of severe burns better recapitulates standard clinical care, some limitations in this model remain. After wound excision, open wounds are typically covered with an occlusive wound dressing or donor skin. While another study utilizing a rodent burn+excision model included subsequent wound dressings^25^, we did not cover wounds with occlusive dressings or skin grafts in our current study. With that said, Hew and colleagues performed wound excisions in mice and subsequently covered excised wou nds with dressings, and still observed severe prolonged hypermetabolism despite the use of occlusive burn wound dressing, comparable to that seen in the current study. Further, our approach of acclimating and housing mice at thermoneutrality during their recovery likely reduces evaporative heat loss from open wounds.

Collectively, by leveraging high-resolution physiological phenotyping and thermoneutral housing to validate and refine a mouse model of scald burns, we demonstrate that wound excision is a primary driver of acute hypermetabolism and cachexia post-injury. Accordingly, our data suggest that in order to accurately model the acute hypermetabolic and cachectic responses to burns and, thus, to promote rigor, reproducibility and the translational value mouse models of burn trauma, robust metabolic phenotyping and burn wound excision should be incorporated into these preclinical models.

## Supplemental Figures

**Supplemental Figure 1:**
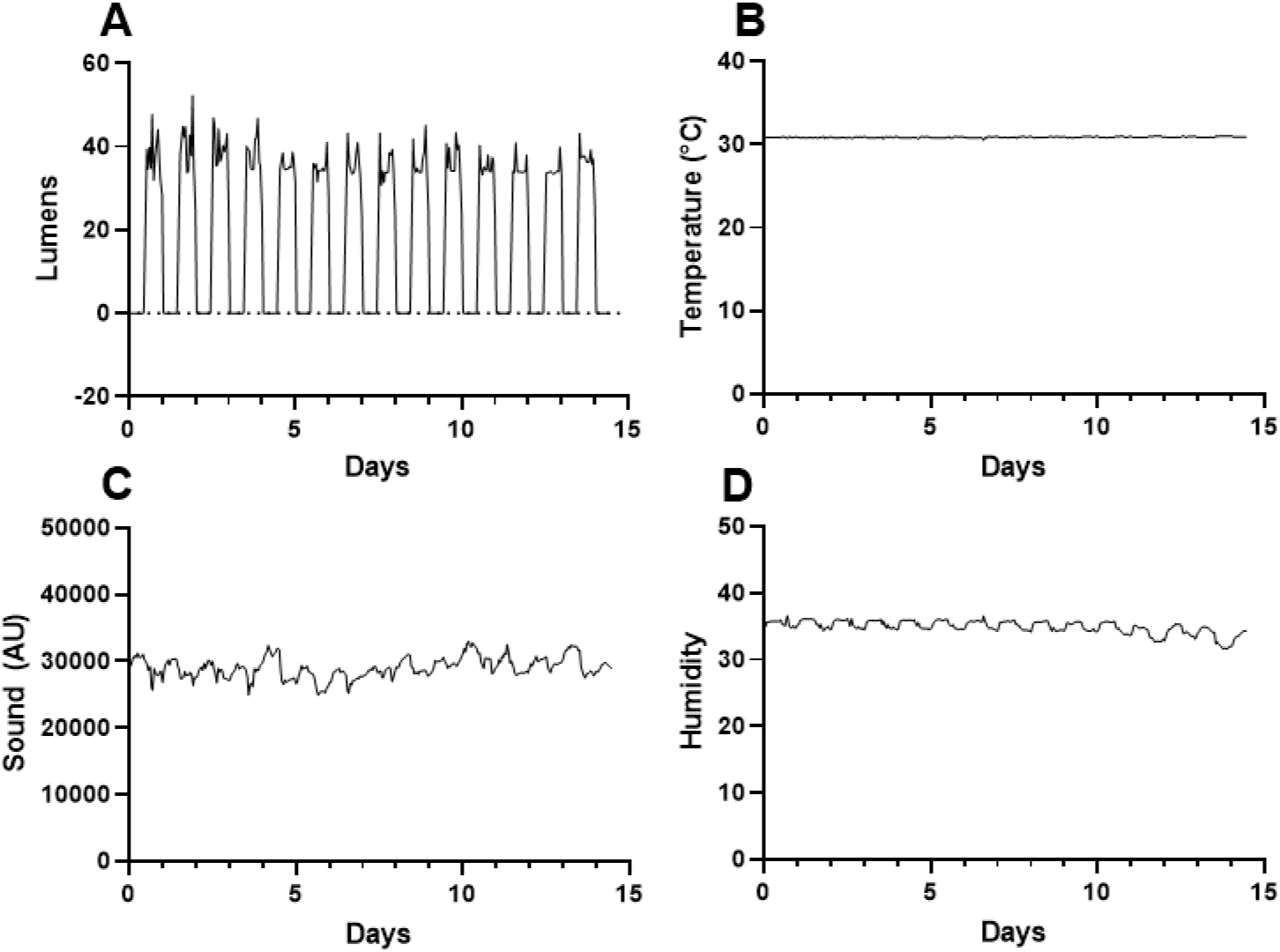
Light Intensity (A), temperature (B), ambient sound (C), and humidity (D) measured throughout the course of the study.

**Supplemental Figure 2:**
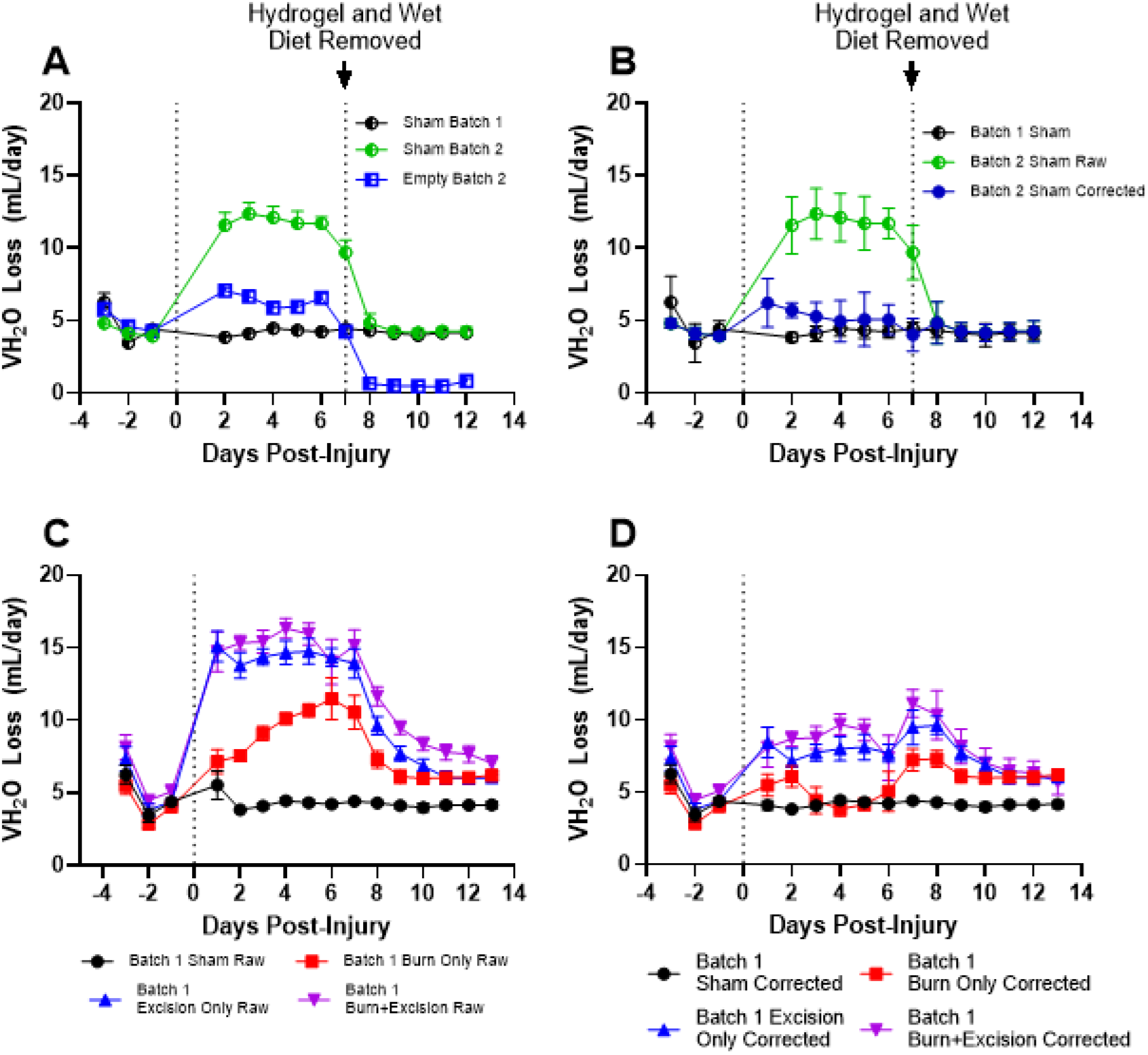
The daily evaporative water loss in sham mice in batch 1 (no hydrogel/wet diet), sham mice in batch 2 (hydrogel/wet diet for 7 days post-injury), and empty cages (hydrogel/wet diet for 7 days post-injury) (A). The daily evaporative water loss in sham mice in batch 1, sham mice in batch, and empty cages corrected for the evaporative water loss caused by supplemental hydrogel/wet diet (B). The raw daily evaporative water loss in sham mice (no hydrogel/wet diet) and burn only, excision only, burn + excision mi ce in batch 1 given hydrogel/wet diet for 7 days post-injury (C). The daily evaporative water loss in batch 1 mice corrected for evaporative water loss by hydrogel/wet diet (D).

**Supplemental Figure 3:**
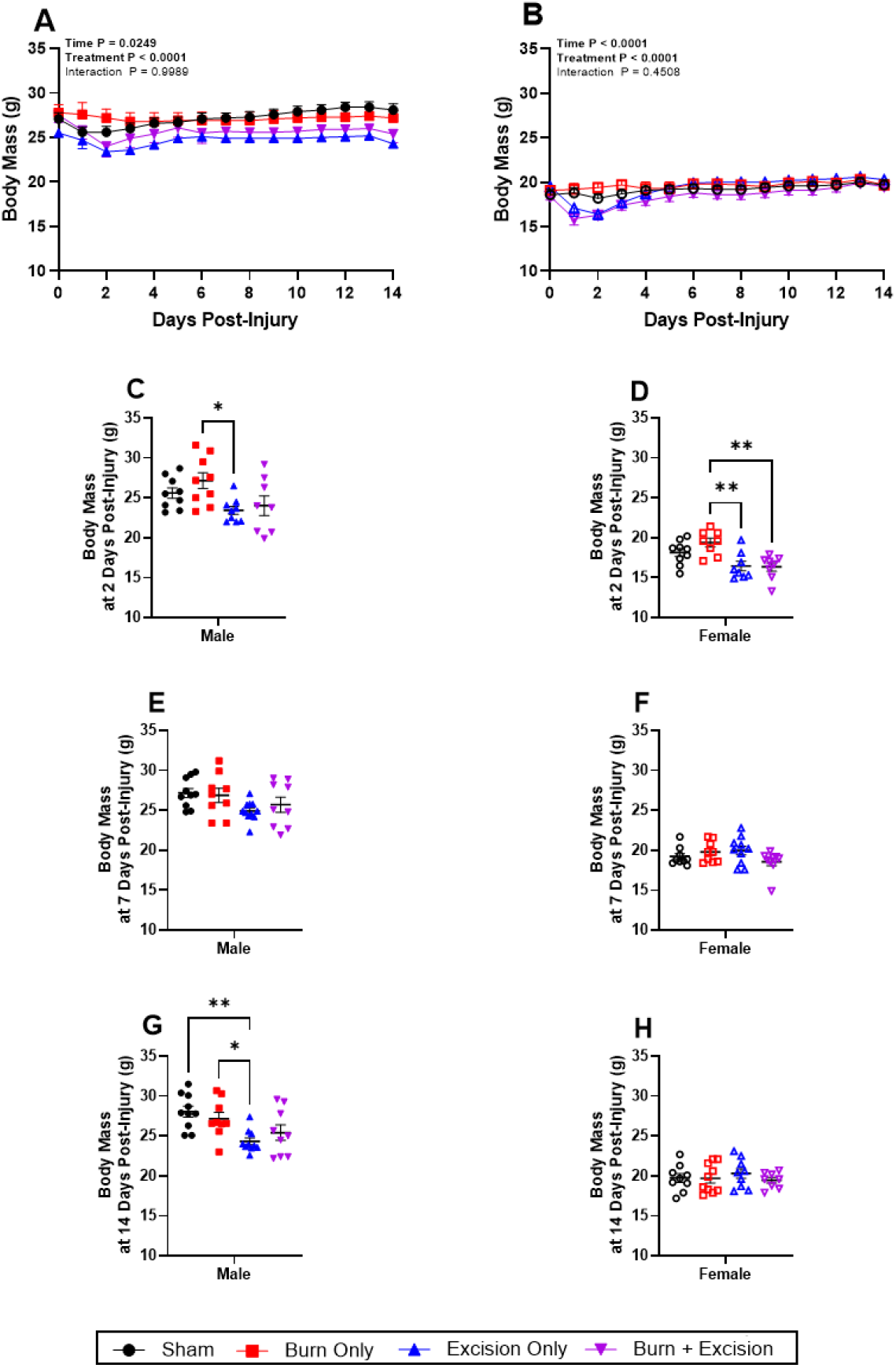
Change from in body mass in male (A) and female (B) mice. Data show percent change in body mass from day 0 prior to procedure and during the 14-day recovery period. Body mass is also shown at days 2, 7, and 14 post-injury in male (panels C, E, and G) and female (panels D, F, and H) mice. Values are means±SEM, n=9-11 per group. Panels A and B analyzed using Two-Way ANOVAS, with Tukey’s test for Multiple Comparisons comparing between groups over time. On all other panels, One-Way ANOVAS, with Tukey’s test for Multiple Comparisons were used to compare groups. *P<0.05, **P<0.01, ***P<0.001 and ****P<0.0001.

**Supplemental Figure 4:**
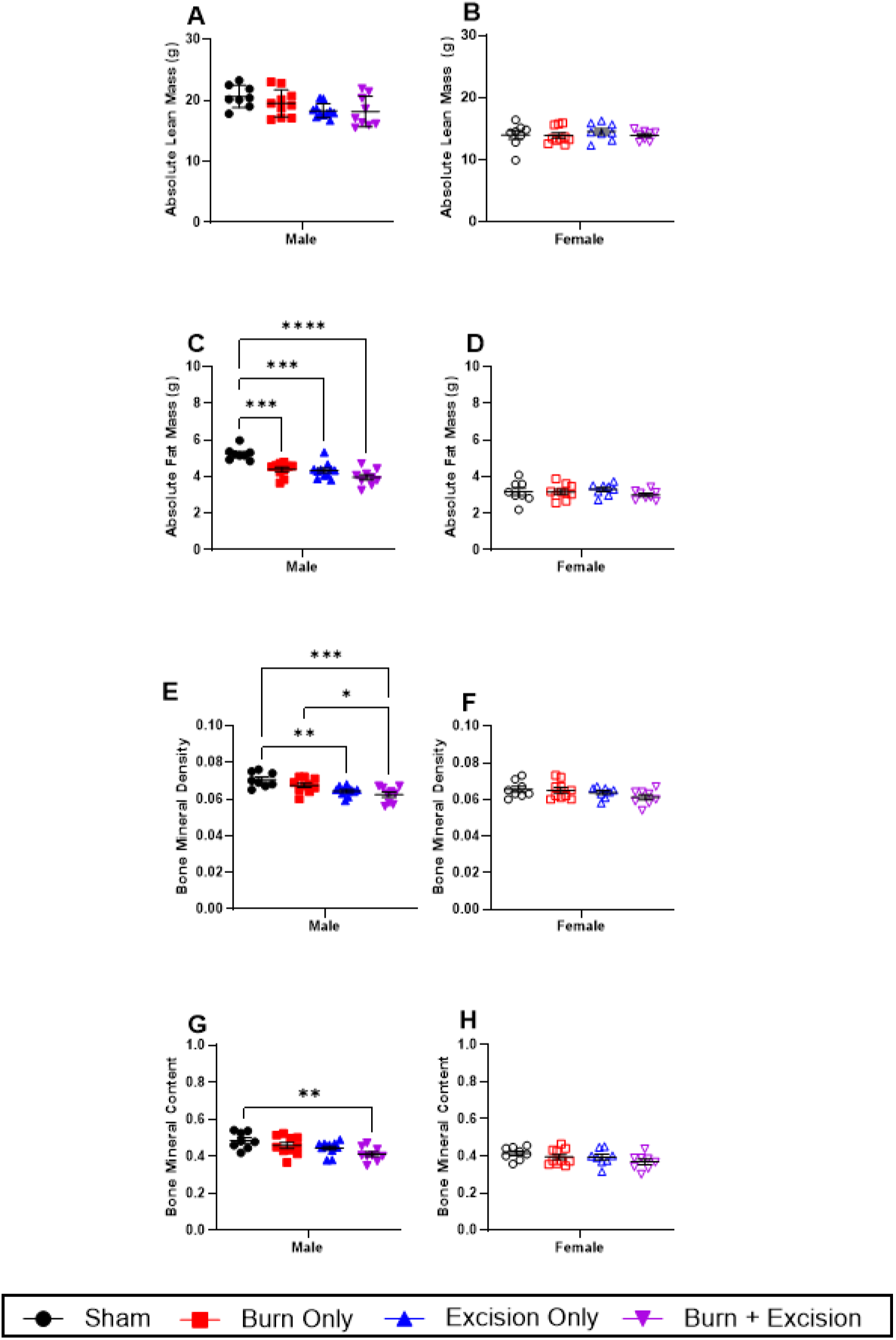
Absolute lean mass in male (A) and female (B) mice. Absolute fat mass in male (C) and female (D) mice. Bone mineral density in male (E) and female (F) mice. Bone mineral content in male (G) and female (H) mice Values are means±SEM, n=9-11 per group. One-Way ANOVAS, with Tukey’s test for Multiple Comparisons were used to compare groups. *P<0.05, **P<0.01, ***P<0.001 and ****P<0.0001.

**Supplemental Figure 5:**
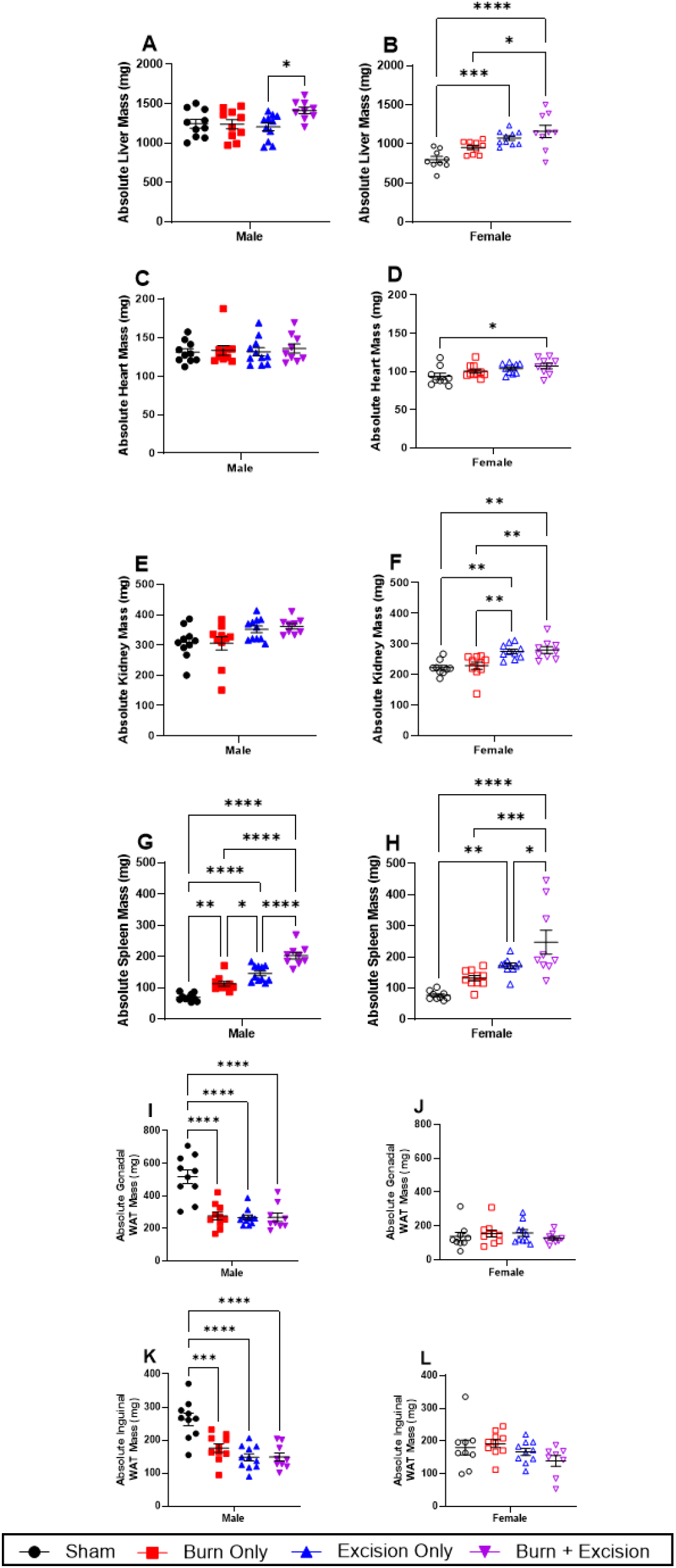
Liver size in male (A) and female (B) mice. Heart size in male (C) and female (D) mice. The kidneys size in male (E) and female (F) mice. The spleen size in male (G) and female (H) mice. The gonadal white adipose tissue depot size in male (I) and female (J) mice. The inguinal white adipose tissue depot size in male (K) and female (L) mice. All organs taken at day 14 immediately after euthanasia. Values are means ± standard errors, n=9-11 per group. One-Way ANOVAS, with Tukey’s test for Multiple Comparisons were used to compare groups. *P<0.05, **P<0.01, ***P<0.001 and ****P<0.0001.

